# Development of human lateral prefrontal sulcal morphology and its relation to reasoning performance

**DOI:** 10.1101/2022.09.14.507822

**Authors:** Ethan H. Willbrand, Emilio Ferrer, Silvia A. Bunge, Kevin S. Weiner

## Abstract

Previous findings show that the morphology of folds (sulci) of the human cerebral cortex flatten during postnatal development. However, previous studies did not consider the relationship between sulcal morphology and cognitive development in individual participants. Here, we fill this gap in knowledge by leveraging cross-sectional morphological neuroimaging data in the lateral prefrontal cortex (LPFC) from individual human participants (6-36 years old, males and females; N = 108; 3672 sulci), as well as longitudinal morphological and behavioral data from a subset of child and adolescent participants scanned at two timepoints (6-18 years old; N = 44; 2992 sulci). Manually defining thousands of sulci revealed that LPFC sulcal morphology (depth, surface area, gray matter thickness, and local gyrification index) differed between children (6-11 years old)/adolescents (11-18 years old) and young adults (22-36 years old) cross-sectionally, but only cortical thickness showed both cross-sectional differences between children and adolescents and presented longitudinal changes during childhood and adolescence. Furthermore, a data-driven approach relating morphology and cognition identified that longitudinal changes in cortical thickness of four rostral LPFC sulci predicted longitudinal changes in reasoning performance, a higher-level cognitive ability that relies on LPFC. Contrary to previous findings, these results suggest that sulci may flatten either after this time frame or over a longer longitudinal period of time than previously presented. Crucially, these results also suggest that longitudinal changes in the cortex within specific LPFC sulci are behaviorally meaningful—providing targeted structures, and areas of the cortex, for future neuroimaging studies examining the development of cognitive abilities.

**Significance Statement:** Recent work has shown that individual differences in neuroanatomical structures (indentations, or sulci) within the lateral prefrontal cortex (LPFC) are behaviorally meaningful during childhood and adolescence. Here, we describe how specific LPFC sulci develop at the level of individual participants for the first time—from both cross-sectional and longitudinal perspectives. Further, we show, also for the first time, that the longitudinal morphological changes in these structures are behaviorally relevant. These findings lay the foundation for a future avenue to precisely study the development of the cortex and highlight the importance of studying the development of sulci in other cortical expanses and charting how these changes relate to the cognitive abilities those areas support at the level of individual participants.

## Introduction

Over the past decades, progress has been made toward understanding how the anatomy of the cerebral cortex broadly changes during human development. For example, prior work has shown that cortical gray matter decreases while cortical white matter and surface area increase (Gogtay et al., 2004; Sowell et al., 2004; Lebel et al., 2008; Shaw et al., 2008; Brown et al., 2012; Amlien et al., 2016; Tamnes et al., 2017; de Faria et al., 2021; Norbom et al., 2021; Baum et al., 2022; Bethlehem et al., 2022; Fuhrmann et al., 2022). Nevertheless, a critical gap in knowledge remains: How do neuroanatomical structures within the cerebral cortex—especially those that are hominoid-specific—develop at the individual level?

To address this question, the present study focuses on the development of lateral prefrontal cortex (LPFC) for several key reasons. First, LPFC is relatively enlarged in humans compared to other commonly studied nonhuman primates (Semendeferi et al., 2002; Donahue et al., 2018). Second and third, LPFC is one of the slowest maturing cortical regions (Gogtay et al., 2004; Tamnes et al., 2013; Chini and Hanganu-Opatz, 2021) and plays a central role in many later-developing cognitive abilities such as reasoning (Luria, 1966; Milner and Petrides, 1984; Krawczyk, 2012; Stuss and Knight, 2013). Fourth, LPFC contains indentations, or sulci, that are either hominoid-specific or human-specific (Amiez and Petrides, 2007; Petrides, 2019; Miller et al., 2021a, 2021b; Voorhies et al., 2021; Hathaway et al., 2022; Willbrand et al., 2022b; Yao et al., 2022). Importantly, the morphology (Voorhies et al., 2021; Yao et al., 2022) and the presence or absence of these variable LPFC sulci (Willbrand et al., 2022b) is related to cognitive abilities. Thus, human LPFC is a heterogeneous entity with sulci that show striking individual differences in morphology that are related to individual differences in cognition—a relationship that necessitates studying LPFC sulcal development at the individual level.

Indeed, analyses at the lobular or group level would likely fail to accurately capture the developmental trajectories of these cortical structures because such analyses obscure this individual variability as previously shown (Miller et al., 2021a, 2021b; Voorhies et al., 2021). For example, one study showed that the cortex “flattens” at the lobular level across ages 11-17, exhibiting increases in sulcal width and decreases in sulcal depth, thickness, and local gyrification—especially in the frontal lobe (Alemán-Gómez et al., 2013). Nevertheless, while these results have helped guide the field, it is presently unknown if these findings extend to analyses conducted at a more granular level—such as the development of LPFC sulci at the individual level. Accordingly, age-related changes in the majority of these LPFC sulci and the behavioral relevance of such changes are largely unknown.

Given the limitations of “large N,” group-level analyses on studying sulcal morphology (Miller et al., 2021a, 2021b), here, we took a “deep imaging” approach (Gratton et al., 2022) to study age-related changes in sulcal morphology and brain-behavior relationships by focusing on a large sampling of individual sulci within individual participants. To this end, we first leveraged individual LPFC sulcal data (3672 sulcal definitions) from 108 participants (6-36 years old) from previous work in pediatric (Voorhies et al., 2021; Willbrand et al., 2022b; Yao et al., 2022) and young adult samples (Miller et al., 2021b) to address whether the morphology of these sulci differs cross-sectionally during this timeframe. We then defined 2992 sulci (1496 per timepoint) in a subset of the pediatric participants scanned at two timepoints (N = 44) to determine whether LPFC sulcal morphology changes longitudinally, across an average timeframe of 1.6 years. Finally, we tested whether longitudinal changes in LPFC sulcal morphology predicted longitudinal changes in a reasoning task that relies on LPFC (Milner and Petrides, 1984; Christoff et al., 2001; Crone et al., 2009; Vendetti and Bunge, 2014).

## Materials and Methods

### Participants

#### Children and adolescents

##### General details

The present study leveraged previously published data from the longitudinal Neurodevelopment of Reasoning Ability (NORA) dataset (Wendelken et al., 2011, 2016, 2017; Ferrer et al., 2013; Voorhies et al., 2021; Willbrand et al., 2022b; Yao et al., 2022). All participants were screened for neurological impairments, psychiatric illness, history of learning disability, and developmental delay. Participants and their parents gave their informed assent and/or consent to participate in the study, which was approved by the Committee for the Protection of Human Participants at the University of California, Berkeley.

##### Cross-sectional sample

72 typically developing individuals (30 females and 42 males, based on parent-reported gender identity) between the ages of 6-18 (average age ± sd = 12.06 ± 3.69 years) were randomly selected from the dataset. The descriptive statistics represent the ages of the selected participants at the time of the selected scan. These participants were the same as those used in previous studies examining the relationship between the morphology of LPFC sulci and behavior (Voorhies et al., 2021; Willbrand et al., 2022b; Yao et al., 2022). For cross-sectional analyses, we divided this pediatric sample along the median age (11.6 years) to establish two equally sized age groups (*children*: mean age ± sd = 8.90 ± 1.59 years, 6.41-11.53 years old; *adolescents*: mean age ± sd = 15.21 ± 2.15 years, 11.66-18.86 years old) to the young adult sample (**Table 1**). Additional demographic and socioeconomic data are included in Extended Data **Table 1-1**.

**Table 1.** Gender and age demographics of the three age groups. The age column depicts age ± sd. Additional demographic information is included in Extended Data Tables 1-1 and 1-2.

| Age group | n (%F) | Age (years) | Age range (years) |
| --- | --- | --- | --- |
| Children | 36 (47.2) | 8.9 ± 1.59 | 6.41-11.53 |
| Adolescents | 36 (36.1) | 15.21 ± 2.15 | 11.66-18.86 |
| Young Adults | 36 (47.2) | 28.97 ± 3.78 | 22-36 |

##### Longitudinal sample

Of the 72 children and adolescents in the cross-sectional analyses, 49 participants (19 females and 30 males) were scanned at two separate timepoints (average age at baseline ± sd = 10.82 ± 3.44 years, 6.25-17.65; average age at follow-up ± sd = 12.37 ± 3.58 years, 7.57-19.32 years; average number of years between scans ± sd = 1.55 ± 0.44 years). These initial 49 participants were tentatively selected for longitudinal morphological analyses; however, due to subsequent selection criteria (see below), such morphological analyses were conducted on 44 of these participants (females = 18, males = 26; average age at baseline ± sd = 11.1 ± 3.47 years; average number of years at follow-up ± sd = 12.66 ± 3.61 years; average number of years between scans ± sd = 1.56 ± 0.46 years).

#### Young adults

We analyzed young adult structural MRI data from the freely available Human Connectome Project database (HCP; https://www.humanconnectome.org/study/hcp-young-adult) (Van Essen et al., 2012). This cross-sectional dataset consisted of 36 randomly selected HCP participants (17 females and 19 males) whose ages were between 22 and 36 (average age ± sd = 28.97 ± 3.78 years). See Extended Data **Table 1-2** for additional demographic and socioeconomic information. These participants were the same as those used in a previous study examining the anatomical and functional features of LPFC sulci (Miller et al., 2021b). These data were previously acquired using protocols approved by the Washington University Institutional Review Board.

### Imaging data acquisition

#### Children and adolescents

High-resolution T1-weighted MPRAGE anatomical scans (TR = 2300 ms, TE = 2.98 ms, 1 mm isotropic voxels) were acquired using the Siemens 3T Trio fMRI scanner at the University of California Berkeley Brain Imaging Center. For participants included in the longitudinal analyses, two anatomical scans were collected across an average delay of 1.56 ± 0.46 years (0.87-3.28 years) between timepoints.

#### Young adults

Anatomical T1-weighted MPRAGE anatomical scans (0.8 mm isotropic voxel resolution) were obtained in native space from the HCP database, along with outputs from the HCP modified FreeSurfer pipeline (Dale et al., 1999; Fischl et al., 1999a; Glasser et al., 2013). Additional details on image acquisition parameters and image processing can be found in (Glasser et al., 2013).

### Morphological analyses

#### Cross-sectional cortical surface reconstruction pipeline

Each T1-weighted image was visually inspected for scanner artifacts. Afterward, reconstructions of the cortical surfaces were generated for each participant from their T1 scans using a standard FreeSurfer pipeline (FreeSurfer (v6.0.0): surfer.nmr.mgh.harvard.edu/) (Dale et al., 1999; Fischl et al., 1999a, 1999b). Cortical reconstructions were created from the resulting boundary made by segmenting the gray and white matter in each anatomical volume with FreeSurfer’s automated segmentation tools (Dale et al., 1999). Each reconstruction was inspected for segmentation errors, which were then manually corrected when necessary. Subsequent sulcal labeling and extraction of anatomical metrics were calculated from individual participants’ cortical surface reconstructions.

#### Longitudinal cortical surface reconstruction pipeline

The FreeSurfer longitudinal Stream (v6.0.0; https://surfer.nmr.mgh.harvard.edu/fswiki/LongitudinalProcessing) (Reuter et al., 2012) was leveraged to create accurate and unbiased cortical reconstructions of the two independent scans that were obtained for each individual participant for whom we had scans at two timepoints. Here, we briefly describe this longitudinal processing pipeline (see (Reuter et al., 2010, 2012; Reuter and Fischl, 2011) for additional details). First, the two T1-weighted images for each participant are processed independently using the standard FreeSurfer pipeline described previously (Dale et al., 1999; Fischl et al., 1999a, 1999b). Second, an unbiased within-subject template space and image (*base template*) is created from these two independent images using robust, inverse consistent registration (Reuter et al., 2010; Reuter and Fischl, 2011). Third, after the creation of this base template, these independent, cross-sectional images are processed again; however, several processing steps (e.g., skull stripping, Talairach transforms, atlas registration as well as spherical surface maps and parcellations) are initialized with common information from the within-subject template. This procedure significantly increases reliability and statistical power, as well as the robustness and sensitivity of the overall longitudinal analysis (Reuter et al., 2010, 2012). In addition, new probabilistic methods were applied to further reduce the variability of results across timepoints.

Each reconstruction was also inspected for major topological errors between the first two steps. Participants were excluded if one of their reconstructions was of insufficient quality. In total, five participants (females = 1, males = 4; average age at baseline ± sd = 8.4 ± 2.18 years; average age at follow-up ± sd = 9.82 ± 2.22 years; average number of years between scans ± sd = 1.43 ± 0.15 years) were excluded due to having a poor reconstruction at one timepoint, resulting in the final sample of 44 participants for longitudinal morphological analyses.

#### General procedure for manual labeling of LPFC sulci

The majority of the sulci described here were originally defined in these two samples for previous work studying the relationship between LPFC sulci and anatomical, functional, and behavioral features (see Miller et al., 2021a, 2021b; Voorhies et al., 2021; Willbrand et al., 2022b; Yao et al., 2022). Sulcal labels in the LPFC were based on the most recent parcellation proposed by Petrides (2019; see also Sprung-Much and Petrides, 2018, 2020).

LPFC sulci were manually defined in each hemisphere and each participant on both the pial and inflated surfaces (for further details refer to Miller et al., 2021b; Voorhies et al., 2021). An example hemisphere is shown in **Figure 1A**, *left* (see Extended Data **Figure 1-1** for a summary of the 18 identifiable LPFC sulci). We first defined the three larger sulci that serve as the posterior boundary of LPFC, namely the 1) *central sulcus* (cs), as well as 2) the *superior* (sprs) and 3) *inferior* (iprs) components of the *precentral sulcus*. We then defined the large sulcus running longitudinally across LPFC: the *inferior frontal sulcus* (ifs). We then defined eight sulci anterior to the prcs and superior to the ifs: 1) the *anterior* (sfs-a) and 2) *posterior* (sfs-p) components of the *superior frontal sulcus* (sfs), 3) the *vertical* (imfs-v) and 4) *horizontal* (imfs-h) components of the *intermediate fronto-marginal sulcus* (imfs), 5) the *anterior* (pmfs-a), 6) *intermediate* (pmfs-i), and 7) *posterior* (pmfs-p) components of the *posterior middle frontal sulcus* (pmfs), and 8) the *para-intermediate frontal sulcus* (pimfs). These 12 sulci have previously been identified in children and adolescents (Voorhies et al., 2021; Willbrand et al., 2022b; Yao et al., 2022), as well as in young adults (Miller et al., 2021b). Finally, we defined the six sulci inferior to the *ifs*: the *diagonal sulcus* (ds), the *triangular sulcus* (ts), the *pre-triangular sulcus* (prts), the *lateral marginal frontal sulcus* (lfms), the *ascending ramus of the lateral fissure* (aalf), and the *horizontal ramus of the lateral fissure* (half). These six sulci were previously identified in children and adolescents by (Yao et al., 2022), and in young adults for the present study. One of these 18 sulci, *pimfs*, showed a high degree of variability, with individuals exhibiting 2, 1, or 0 branches in a given hemisphere (Voorhies et al., 2021; Willbrand et al., 2022b); as a result, this sulcus was not included in our developmental morphological analyses. All in all, the present study considered 17 LPFC sulci that were reliably identified across participants and hemispheres.

**Figure 1.**
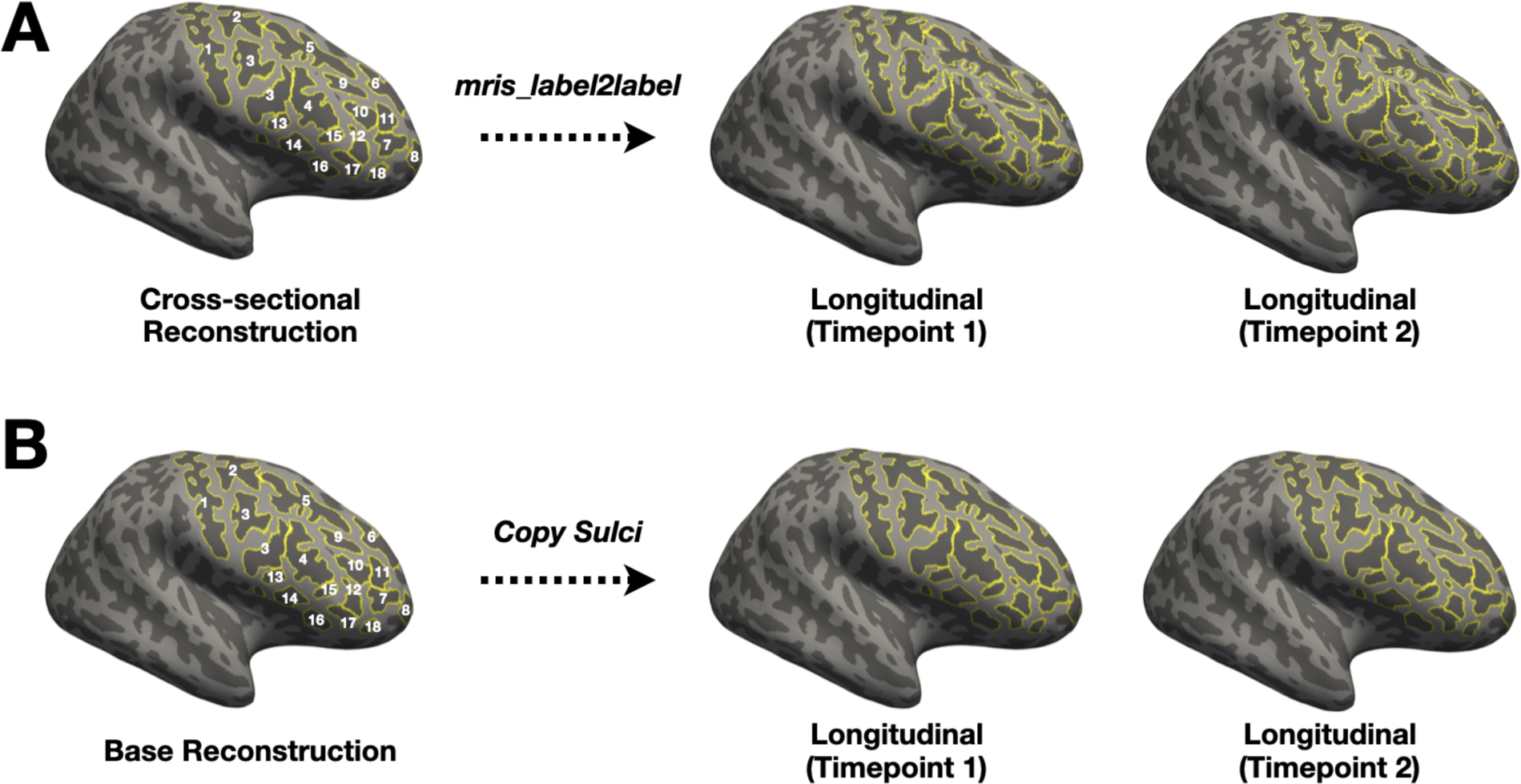
Pipeline for defining individual LPFC sulci longitudinally. ***A***. Cross-sectional (*left*) and longitudinal (*right*) inflated cortical surfaces (sulci: dark gray; gyri: light gray) for an example participant. LPFC sulci are outlined in yellow and the numbers identifying each sulcus correspond with those in Extended Data Figure 1-1. Copying sulcal labels from the cross-sectional surfaces (Materials and Methods) to the longitudinal surfaces with *mris_label2label* does not provide accurate sulcal definitions. ***B***. The same format as *A*, except with the base template (*left*) inflated cortical surface for this example participant. For the longitudinal analyses, LPFC sulci were redefined on the base surfaces (Materials and Methods) since copying these labels from the base template to the longitudinal surfaces provides highly accurate individual labels on the longitudinal surfaces—in contrast to *A*. This process ensures that each sulcus is indeed the same structure at both time points.

As in our previous work, each LPFC sulcus was manually defined within each individual hemisphere on the FreeSurfer *inflated* mesh with tools in *tksurfer*, which enables accurate definition of individual sulci within *in vivo* MRI data (Weiner et al., 2014, 2018; Miller et al., 2021a, 2021b; Voorhies et al., 2021). Specifically, the *curvature* metric in FreeSurfer distinguished the boundaries between sulcal and gyral components. Manual lines were drawn on the inflated cortical surface to define sulci based on the proposal by (Petrides, 2019), and guided by the *pial* and *smoothwm* surfaces of each individual. Since the precise start or end point of a sulcus can be difficult to determine on one surface (Borne et al., 2020), using the *inflated*, *pial*, and *smoothwm* surfaces of each individual allows us to form a consensus across surfaces to clearly determine each sulcal boundary. The sulcal labels were generated using a two-tiered procedure. The location and definition of each label were first confirmed through a qualitative inter-rater reliability process (E.H.W., W.I.V., J.K.Y., I.R., and J.A.M.) and then finalized by a neuroanatomist (K.S.W.). All anatomical labels for a given hemisphere were fully defined before any morphological analyses of the sulcal labels were performed.

#### Modified procedure to manually label LPFC sulci longitudinally

After obtaining the two longitudinal reconstructions for the 44 participants, we first redefined the LPFC sulci on the *base* template of each participant (see **Fig. 1B**, *left*, for an example hemisphere). This first step was performed since defining sulci on the base template surface and copying these labels to the longitudinal surfaces provides far more accurate sulcal definitions compared to using the *mris_label2label* function to convert labels from the cross-sectional surface to the longitudinal surfaces (**Fig. 1**). Once all sulci were defined on the base template, the sulcal labels were copied to the two timepoints to extract sulcal morphological features at each timepoint. When necessary, minor corrections were made to sulcal labels at each timepoint before extracting morphological features. The sulcal definitions and corrections in this step were completed by E.H.W. and K.S.W.

#### Extracting morphological features from sulcal labels

After defining LPFC sulci in FreeSurfer, we extracted the following morphological metrics from each sulcal label file: sulcal depth (mm), surface area (mm^2^), cortical thickness (mm), and local gyrification index (lGI). We selected sulcal depth and surface area as these metrics are prominent features of sulci (Sanides, 1964; Chi et al., 1977; Welker, 1990; Armstrong et al., 1995; Weiner et al., 2014, 2018; Lopez-Persem et al., 2019; Madan, 2019; Petrides, 2019; Weiner, 2019; Miller et al., 2020, 2021a, 2021b; Natu et al., 2021; Voorhies et al., 2021; Li et al., 2022; Willbrand et al., 2022a, 2022b; Yao et al., 2022). We further selected cortical thickness and lGI since the longitudinal development of these features was previously analyzed at the lobular level (Alemán-Gómez et al., 2013).

Raw depth values (in mm) were calculated using a custom-modified version of a recently developed algorithm built on the FreeSurfer pipeline (Madan, 2019). Briefly, depth was calculated as the distance between the sulcal fundus and the smoothed outer pial surface (Madan, 2019). Mean thickness and surface area values were extracted from each sulcus using the built-in *mris_anatomical_stats* function in FreeSurfer (Fischl and Dale, 2000). Mean lGI values (a value comparing the amount of cortex buried within sulci to the amount of cortex on the outer visible cortex, or hull) were extracted with the *mris_anatomical_stats* function with the *pial_lgi* flag in FreeSurfer (Fischl and Dale, 2000). As in prior work (Miller et al., 2020; Voorhies et al., 2021; Willbrand et al., 2022a, 2022b), we also calculated normalized values of thickness, depth, and surface area values—to the thickest and deepest points, and total surface area of the cortex, respectively. Consistent with prior studies (Alemán-Gómez et al., 2013; Klein et al., 2014; Cao et al., 2017; Forde et al., 2017; Libero et al., 2019; Gharehgazlou et al., 2021), we did not normalize lGI values.

Although prior work examining sulcal morphology studied sulcal width (for example see Alemán-Gómez et al., 2013; Jin et al., 2018; Carmona et al., 2019), we found that the current toolbox that can extract this information from individual sulcal labels in FreeSurfer (Madan, 2019) failed to obtain the width of many of the smaller sulci in LPFC. Thus, we were not able to examine this feature in the present study. With improved methodology, future work should seek to examine how the width of all LPFC sulci changes during development.

#### Classifying LPFC sulci based on morphology

All morphological analyses were implemented in R (v4.1.2; http://www.r-project.org). We used an unsupervised, data-driven approach to group LPFC sulci based on their morphology: namely, a combination of k-means clustering and multi-dimensional scaling (MDS) using the four morphological features of sulci analyzed in this study (sulcal depth, surface area, cortical thickness, and lGI). These analyses were run on scaled morphological data, separately in each hemisphere. Importantly, these analyses were conducted in a manner that was blind to age group (children, adolescents, and young adults) and sulcal type (primary, secondary, and tertiary; see Miller et al. 2021a, 2021b; Voorhies et al., 2021; Yao et al., 2022 for further details). Further, rather than choosing the number of clusters (k) ourselves or with a single index, we quantitatively determined the optimal number of clusters using the *NbClust* function (from the *NbClust* R package). Briefly, this function leverages 30 indices to propose the best number of clusters for the data based on the majority rule (for additional details see Charrad et al., 2014).

In both hemispheres, the majority of indices determined that three was the optimal number of clusters to describe the data. K-means clustering was then implemented using the *kmeans* function (from the *stats* R package) with the optimal number of clusters (k = 3) suggested by the *NbClust* function. Next, given the multidimensionality of the dataset (dimensions = 4), we ran classic (metric) MDS with the *dist* and *cmdscale* R functions (from the *stats* R package) to visualize the (dis)similarity of the data and clusters in a lower-dimensional space (dimensions = 2). Cluster plots were then generated (using the *ggscatter* function from the *ggpubr* R package) to visualize the results. Finally, for subsequent morphological analyses, the assignment of each LPFC sulcus to a cluster was determined by the majority rule—that is, it was based on what cluster the majority of participants fell into for each sulcus.

#### Comparing the morphology of LPFC sulci between age groups cross-sectionally

To compare the depth, thickness, surface area, and lGI of LPFC sulci between the three established age groups (children, adolescents, and young adults; **Table 1**), we used mixed-model analyses of variance (ANOVAs). For each ANOVA, *hemisphere* (left, right) and *sulcal cluster* (3 clusters) were used as within-participant factors, while *age group* (children, adolescent, young adult) and *gender* (male, female) were implemented as between-participant factors. *Hemisphere* and *gender* were included due to prior research showing hemispheric differences in sulcal development (Chi et al., 1977) and mixed results regarding gender differences in sulcal development (Welker, 1990; Gottfredson, 1997). The Greenhouse-Geisser sphericity correction method was applied for all ANOVAs, which adjusted the degrees of freedom. The same ANOVAs were repeated with the normalized metrics as the dependent variables. All ANOVAs were performed with the *aov_ez* function (from the *afex* R package). Effect sizes are reported with the generalized eta-squared (η^2^G) metric. Given the large number of tests performed, *p* < .05 was considered significant for any main effect or interaction after controlling for multiple comparisons (via the false discovery rate (FDR)). Post-hoc pairwise comparisons on any significant main effects or interactions were performed with Tukey’s method using the *emmeans* function (from the *emmeans* R package).

#### Assessing whether LPFC sulcal morphology changes longitudinally

Considering the complexity of our longitudinal sulcal data (2 timepoints, 17 LPFC sulci, 2 hemispheres, 3 clusters), for the present study, we averaged values across the sulci within each cluster (**Fig. 2B**) to create three “*cluster averages*” for each morphological feature in each hemisphere for each participant at each timepoint. This grouping was based on the cross-sectional clusters and conducted prior to running longitudinal analyses. We primarily tested for longitudinal changes in cortical thickness, which was the only feature to show differences between children and adolescents in the cross-sectional analyses (see Results). We also regressed cortical thickness on baseline age to test whether this factor influenced the level (the thickness at timepoint 1) and slope (the change from timepoint 1 to timepoint 2) for each cluster. *Gender* was not included in the models since it was not related to either LPFC morphology cross-sectionally (see Results) or changes in cortical thickness for any cluster (*p*s > .41). We also tested sulcal depth given its behavioral (Voorhies et al., 2021; Yao et al., 2022) and developmental (Alemán-Gómez et al., 2013) relevance. Linear growth curve models (LGC) were run to calculate changes in cortical thickness from timepoint 1 to timepoint 2 for each cluster in each hemisphere. LGCs were implemented with the *sem* function (from the *lavaan* R package).

**Figure 2.**
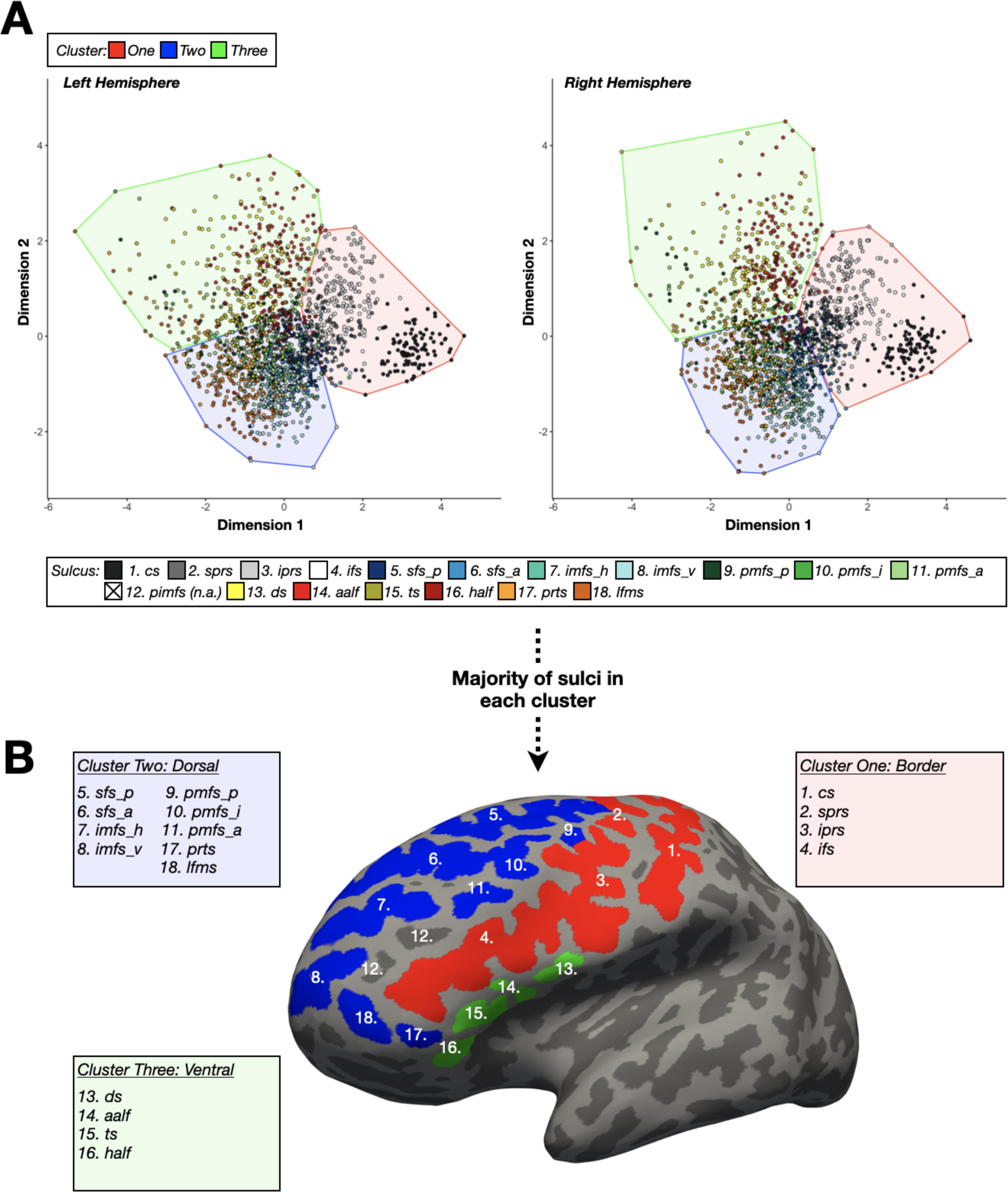
Data-driven approach clusters LPFC sulci into three groups based on morphology. ***A***. Cluster plots visualizing the results of the k-means clustering and multidimensional scaling analysis performed in each hemisphere (*left:* left hemisphere; *right:* right hemisphere). Individual dots represent individual sulci for all 108 participants and are colored by sulcus (see bottom key). The optimal number of clusters that described the LPFC sulci based on morphology was three: one (red), two (blue), and three (green). The sulci in each of the three clusters are encircled by a colored ellipse (see top key). ***B***. Example left hemisphere inflated cortical surface (sulci: dark gray; gyri: light gray). Each LPFC sulcus (excluding the pimfs; Materials and Methods) is colored by the cluster that the majority fell into across participants (as in *A*). The numbers identifying each LPFC sulcus on the cortical surface correspond with those in Extended Data Figure 1-1. Each box contains a list of the abbreviated names of the sulci in each cluster. Generally, Cluster 2 contained dorsal LPFC sulci, whereas Cluster 3 contained ventral LPFC sulci. Cluster 1 contained the sulci that border Clusters 2 and 3 posteriorly or in-between.

### Longitudinal behavioral analyses

#### Behavioral data

The longitudinal pediatric data used here were drawn from a long-running dataset relating cortical anatomy and functional connectivity to reasoning ability (Wendelken et al., 2011, 2016, 2017; Ferrer et al., 2013; Voorhies et al., 2021; Willbrand et al., 2022b; Yao et al., 2022). The majority of participants had scores for the WISC-IV matrix reasoning task (Wechsler, 1949), which is a widely-used, gold-standard measure of abstract, nonverbal reasoning (Ferrer et al., 2013; Wendelken et al., 2016) and is a cognitive ability reliant on LPFC (for review, see Holyoak and Monti, 2021). In the present study, reasoning was specifically measured as a total raw score from the WISC-IV Matrix reasoning task (Wechsler, 1949). Matrix reasoning is an untimed subtest of the WISC-IV in which participants are shown colored matrices with one missing quadrant. The participant is asked to complete the matrix by selecting the appropriate quadrant from an array of options. To maximize our sample size, we focused on the longitudinal relationship between sulcal morphology and matrix reasoning. The behavioral analyses were conducted with the 43 longitudinal pediatric participants (F = 18; Male = 25) who completed the WISC-IV matrix reasoning task at both timepoints (average age at baseline ± sd = 11.1 ± 3.43 years; average number of years at follow-up ± sd = 12.66 ± 3.57 years; average number of years between scans ± sd = 1.56 ± 0.46 years).

#### Model selection

We assessed whether individual differences in changes in sulcal cortical thickness (ΔCT) predicted variability in changes in matrix reasoning performance (ΔMR). As in prior work (Voorhies et al., 2021; Yao et al., 2022), we leveraged a least absolute shrinkage and selection operator (LASSO) regression, separately in each hemisphere, to select which of the LPFC sulci examined in this study, if any, showed ΔCT associated with ΔMR.

A LASSO regression is well-suited to address our question since it facilitates the model selection process and increases the generalizability of a model by providing a sparse solution that reduces coefficient values and decreases variance in the model without increasing bias (Heinze et al., 2018). Further, regularization is recommended in cases where there are many predictors (X > 10), as in this study, because this technique guards against overfitting and increases the likelihood that a model will generalize to other datasets. A LASSO performs L1 regularization by applying a penalty, or shrinking parameter (α), to the absolute magnitude of the coefficients. In this manner, low coefficients are set to zero and eliminated from the model. Therefore, LASSO affords data-driven variable selection that results in simplified models containing only the most predictive features—in this case, sulci predicting reasoning performance. Not only does this methodology improve model interpretability and prediction accuracy, but it also protects against overfitting— thus improving generalizability (Heinze et al., 2018; Ghojogh and Crowley, 2019).

As part of the model selection process, we used cross-validation to optimize the values for α with the GridSearchCV function from the *sklearn* package in Python (https://scikit-learn.org/stable/modules/generated/sklearn.model_selection.GridSearchCV.html). The GridSearchCV function performs an exhaustive search across a range of specified α values. According to the convention (Heinze et al., 2018), we then selected the α value (and corresponding model) that minimized the cross-validated mean-squared error (MSE_cv_).

#### Model comparisons

To specifically characterize the relationship between ΔCT and ΔMR, we compared the model determined by the LASSO regression to two alternative nested models. All models examined only left hemisphere sulci since the LASSO regressions only identified sulci in the left hemisphere as behaviorally relevant (*left hemisphere:* α = 0.07; **Fig. 4A**). The model did not select any right hemisphere LPFC sulci at the α values with the lowest MSE_cv_ (α = 0.5, MSE_cv_ = 27.12). As reasoning performance improves over childhood (Ferrer et al., 2013; Wendelken et al., 2016, 2017), and ΔMR was correlated with baseline age in our sample (r = -0.43, *p* = .004), we included age at baseline as an additional predictor of ΔMR in the three models. Prior to running the three models, we also ensured that individual variability in ΔCT was not reliably related to baseline age (Extended Data **Fig. 4-1**). We did not include another potentially relevant time-related variable (Δtime) in these models as it did not relate to ΔMR (r = -0.04, *p* = .78).

To confirm the results of our selection, we first constructed a simplified linear model with the four left-hemisphere LPFC sulci that were selected by the LASSO regression as the strongest predictors of ΔMR. We refer to this simplified model as the *LASSO-derived model* [1].

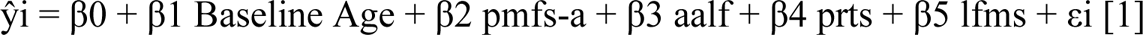

We then compared the fit of the LASSO-derived model with a *full model* that included the ΔCT of all LPFC sulci within the left hemisphere, as well as baseline age [2].

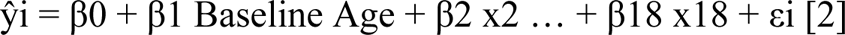

In the nested full model, x2 - x18 represent the change in ΔCT of each of the 17 LPFC sulci in the left hemisphere included in this study, and β2 - β18 represent the associated coefficients.

Finally, we compared our LASSO-derived model, which included both baseline age and the ΔCT of the selected sulci [1], to a model with baseline age as the sole predictor [3]. This nested comparison allowed us to discern whether the LPFC sulci in our selected model explained variance in ΔMR not captured by baseline age alone.

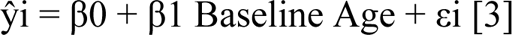

All linear models were fitted with a LOOCV procedure using the SciKit-learn package in Python (Pedregosa et al., 2011). Since these are nested models (wherein the largest model includes all elements in the smaller models), the model with the best fit was determined as the cross-validated model with the lowest MSE_cv_ and the highest R^2^_cv_ values.

### Data availability

The processed data required to perform all statistical analyses and to reproduce all figures for this project will be freely available with the publication of the paper on GitHub (https://github.com/cnl-berkeley/stable_projects). Requests for further information or raw data should be directed to the corresponding authors, K.S.W. and S.A.B.

## Results

### A data-driven approach produced three clusters of LPFC sulci based on a combination of morphological features

Classic and modern studies acknowledge primary, secondary, and tertiary sulci based on several criteria such as the timepoint in which sulci emerge in gestation, depth, surface area, and variability in the presence or absence of sulci. Nevertheless, perhaps unsurprisingly, neuroanatomists have continued to argue about which sulci are lumped into each group for decades. To circumvent these arguments, we implemented a data-driven approach that employed a combination of k-means clustering and multidimensional scaling to test whether the 17 LPFC sulci in the present study could be grouped based on the four morphological features extracted in the present study (Materials and Methods; sulcal depth, surface area, cortical thickness, and lGI).

This analysis revealed that, irrespective of qualitative definitions of primary, secondary, and tertiary sulci, LPFC sulci could be quantitatively grouped into three clusters (**Fig. 2A**). Cluster 1 consisted primarily of *dorsal* PFC sulci (sfs-p, sfs-a, imfs-h, imfs-v, pmfs-p, pmfs-i, pmfs-a, lfms, and prts), Cluster 2 primarily of *ventral* PFC sulci (ts, ds, half, and aalf), and Cluster 3 primarily of PFC sulci *bordering* Cluster 1 and Cluster 2—either in-between the groups (ifs) or posteriorly (cs, sprs, and iprs). **Figure 2B** visualizes these three clusters on an example cortical surface. The border sulci were substantially deeper and larger than both ventral and dorsal sulci (51.5% and 45% deeper on average; 528.3% and 242.9% larger on average; **Fig. 3A-B**). Border and dorsal sulci were thinner than ventral sulci (7.4% and 6.6% on average) and border sulci were slightly thinner than DLFPC sulci (0.8% thinner on average; **Fig. 3C**). Finally, ventral sulci had greater local gyrification index (lGI) values than dorsal and border sulci (41.4% and 17% greater on average), while border sulci had a larger lGI than dorsal sulci (20.9% greater on average; **Fig. 3D**). Given the quantitative determination of these three sulcal clusters, subsequent analyses explored whether the morphology of these sulcal clusters changed i) cross-sectionally between age groups (children ages 6.41-11.53, adolescents ages 11.66-18.86, young adults ages 22-36) and ii) longitudinally in the pediatric sample (ages 6-18 years old).

**Figure 3.**
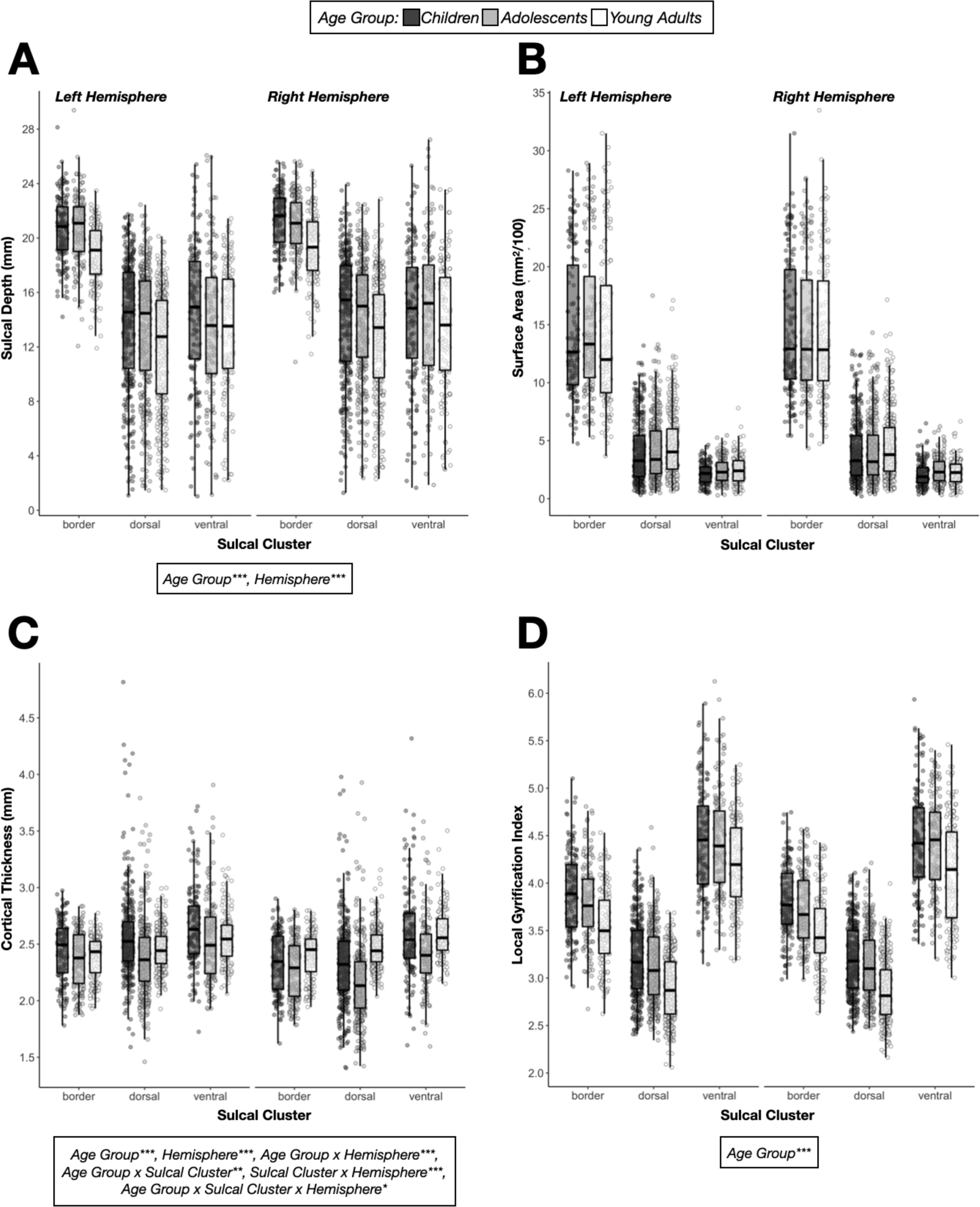
Some, but not all, morphological features of LPFC sulci differ between age groups cross-sectionally. ***A***. Boxplot visualizing sulcal depth (mm) as a function of sulcal cluster (x-axis), age group (colors; black: children, gray: adolescents, white: young adults), and hemisphere (left vs. right faceted plot). Individual dots represent each participant’s individual sulci in each cluster as described in Figure 2B. The text box under the plot displays the significant results from the mixed model ANOVAs shown in Extended Data Figures 3-2 – 3-5 (\**p* < .05; \*\**p* < .01; \*\*\**p* < .001). ***B***. Same as *A,* but for surface area (mm^2^/100). ***C***. Same as *A,* but for mean cortical thickness (mm). ***D***. Same as *A*, but for local gyrification index. These results mostly hold for normalized morphological values (Extended Data Fig. 3-1).

**Figure 4.**
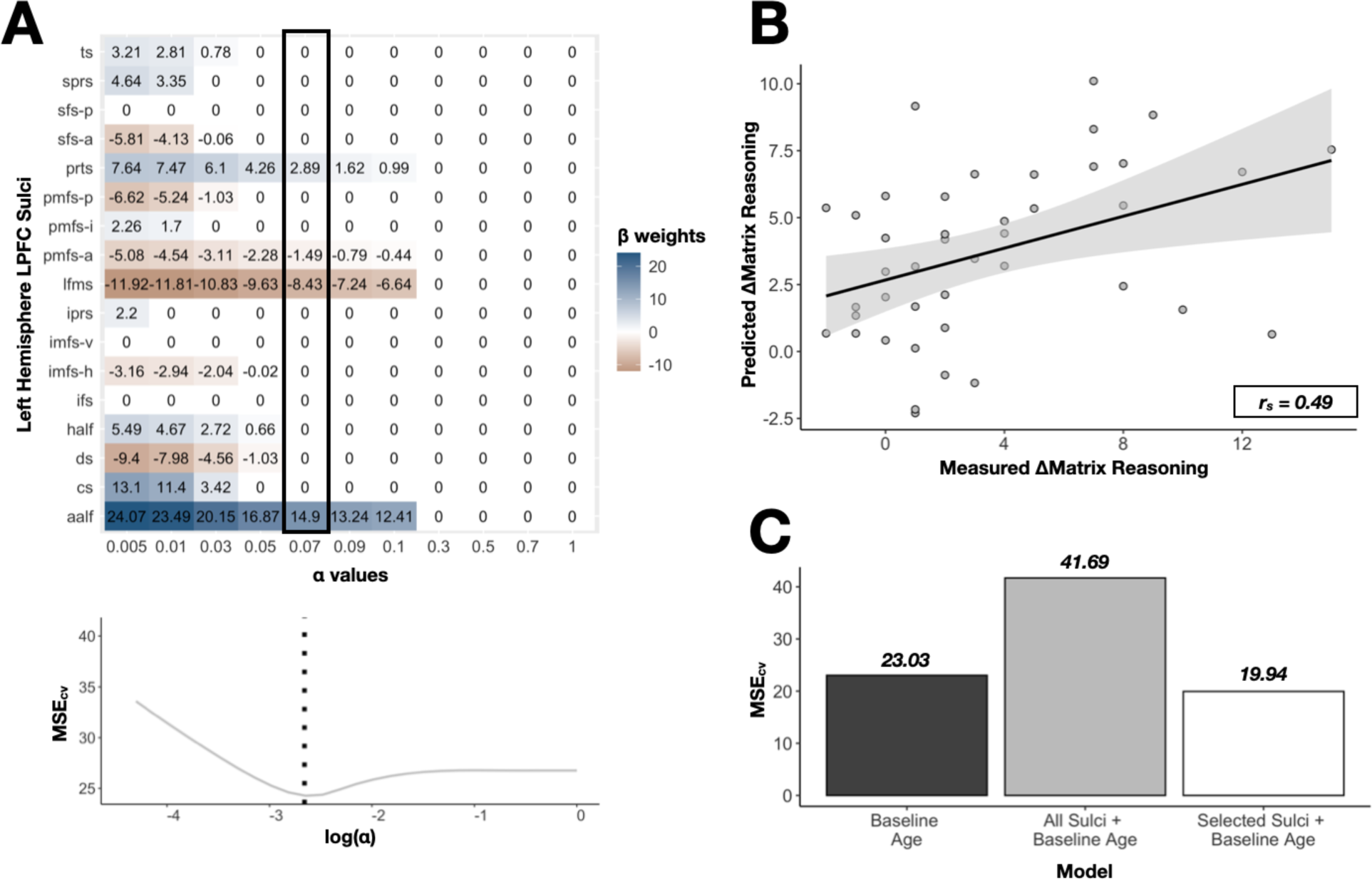
Data-driven model selection reveals that longitudinal changes in cortical thickness of LPFC sulci are associated with longitudinal changes in reasoning performance. Performance and model fits relating change in sulcal cortical thickness (ΔCT) to change in reasoning performance (ΔMR). *A*. *Top*: Beta-coefficients for each sulcus across a range of shrinking parameter (α) values resulting from the LASSO regression in the left hemisphere. The highlighted box indicates the coefficients at the chosen α level with the lowest cross-validated mean-squared error (MSE_cv_). *Bottom*: Cross-validated mean-squared error (MSE_cv_) at each α level. We selected the α value that minimized MSE_cv_ (dotted black line). The LASSO regression with right hemisphere LPFC sulci selected no sulci at the α value that minimized MSE_cv_. *B*. Spearman’s correlation (r_s_ = 0.49) between participants’ measured ΔMR scores and the predicted ΔMR scores from the LOOCV linear regression for the selected model in *A*, which had four left hemisphere LPFC sulci and baseline age as predictors. *D*. Model comparison of the cross-validated MSEs of a model with baseline age as its only predictor (black), a model with all left hemisphere LPFC sulci and baseline age as predictors (gray), and a model with the four selected left hemisphere sulci and baseline age as predictors (white). The model with left- hemisphere sulci selected by the LASSO regression (white) had the lowest MSE_cv_, thus performing the best. Strikingly, the four selected left hemisphere sulci coincide with meta-analysis activation maps for reasoning made with Neurosynth (Fig. 5).

### LPFC sulcal morphology changes cross-sectionally between children, adolescents, and young adults

To quantify cross-sectional differences in LPFC sulcal morphology between age groups, we ran a mixed-model ANOVA for each morphological feature, using within-group factors of *hemisphere* (left, right) and *sulcal cluster* (border sulci, dorsal sulci, ventral sulci) and between-group factors of *age group* (children, adolescents, young adults) and *gender* (male, female). The main results presented below are with the raw morphological metrics unless otherwise specified (see Extended Data **Fig. 3-1** for normalized data and **Figs. 3-2 – 3-5 for** ANOVA results). The only feature to show *gender*-related main effects and interactions was raw surface area; however, these effects were all non-significant with normalized surface area, so we do not report them further (Extended Data **Fig. 3-3**). Finally, we primarily report *age group-*related effects here (see Extended Data **Figs. 3-2 – 3-5** for hemisphere-related effects).

All morphological features displayed a main effect of *age group* except for surface area (Extended Data **Figs. 3-2 – 3-5**). The depth and lGI of LPFC sulci exhibited no differences between adolescents and children (*p*s > .35, Tukey’s adjustment); however, LPFC sulci were deeper and had larger LGI values in children and adolescents than in young adults (*p*s < .0001, Tukey’s adjustment; 11.3% and 9.2% deeper on average, respectively; 9.3% and 7.5% greater lGI on average; **Fig. 3A** and **D**). Conversely, LPFC sulci were thinner in adolescents than in children and young adults (*p*s < .0001, Tukey’s adjustment; 6.1% and 5.7% thinner on average, respectively), but comparable between children and young adults (*p* = .86, Tukey’s adjustment; **Fig. 3C**). There were also complex two- and three-way *age group* interactions with *hemisphere* and *sulcal cluster* for LPFC sulcal thickness, but not for the other three features (**Fig. 3C**; Extended Data **Figs. 3-4** and **3-6**). There were no *age group*-related main effects or interactions (with *sulcal cluster* or *hemisphere*) for raw surface area; however, we observed an *age group* x *sulcal cluster* effect on normalized surface area (see Extended Data **Fig. 3-3**). Broadly speaking, then, we observed decreased sulcal depth and LGI between the pediatric and young adult age groups and a U-shaped pattern for sulcal thickness (thinnest in adolescence). We also observed age-related differences in normalized (but not raw) surface area that differed by sulcal cluster, with dorsal sulci increasing from adolescence to adulthood, ventral sulci increasing incrementally between childhood and adulthood, and border sulci exhibiting no changes across these age groups.

### Cortical thickness of LPFC sulci changes longitudinally during childhood and adolescence

Considering that sulcal cortical thickness was different in adolescents than children cross-sectionally (**Fig. 3C**), we next specifically assessed within-person changes in sulcal cortical thickness during this time period. To this end, we analyzed cortical surfaces for 44 participants in the pediatric sample (6-18 years old) who were scanned at two timepoints (Materials and Methods). LPFC sulci were then defined on the base template and projected to the longitudinal reconstruction at the two points, as this procedure provides an accurate sulcal definition at each timepoint (**Fig. 1**). We then averaged the cortical thickness of the sulci within each cluster (as shown in **Fig. 2B**) at each timepoint. Finally, we implemented linear growth curve models to assess whether the cortical thickness of the LPFC sulcal clusters changed from timepoint 1 to timepoint 2 and whether the baseline age was related to the intercept and the changes.

The cortical thickness of all three LPFC sulcal clusters decreased from timepoint 1 to timepoint 2, to differing degrees. In both hemispheres, the decrease in cortical thickness was greatest for dorsal sulci (*left:* β = -0.078, *z* = -7.71, *p* < .001; *right:* β = -0.098, *z* = -5.76, *p* < .001). In the left hemisphere, ventral sulci showed a comparable decrease in cortical thickness (β = -0.06, *z* = -4.71, *p* < .001) to border sulci (β = -0.059, *z* = -6.87, *p* < .001); in the right hemisphere, on the other hand, ventral sulci showed less of a decrease (β = -0.04, *z* = -2.3, *p* = .021) than border sulci (β = -0.051, *z* = -4.81, *p* < .001). There was no significant variation in the change between participants (*p*s > .22). Further, cortical thickness at timepoint 1 (i.e., each individual’s intercept) differed as a function of baseline age for all clusters (βs ≤ -0.011, *p*s < .001), such that the older the participant, the lower the starting cortical thickness. However, the change in cortical thickness from timepoint 1 to timepoint 2 (i.e., the slope) did not differ by baseline age for any cluster in either hemisphere (-0.006 ≤ βs ≤ 0.001, *p*s > .20). In sum, while all sulcal types showed thinning longitudinally, dorsal sulci showed the most pronounced changes.

Since prior work showed that the average sulcal depth of the frontal lobe declines during adolescence (Alemán-Gómez et al., 2013) and individual differences in sulcal depth are behaviorally-significant (Voorhies et al., 2021; Yao et al., 2022), we also tested whether LPFC sulcal depth changed longitudinally—despite the cross-sectional analysis indicating otherwise (**Fig. 3A**). Indeed, LPFC sulcal depth did not change in our pediatric sample across the average 1.56-year time frame (βs ≤ 0.087, *p*s > .15) and there was no significant variation in the change between participants (*p*s > .21), suggesting that longitudinal changes in LPFC sulcal depth after birth may occur over a longer time span than the one tested here or even after adolescence.

### Longitudinal changes in the cortical thickness of LPFC sulci predict developmental changes in matrix reasoning

Given that LPFC sulcal cortical thickness changed across the two time points, we examined whether these changes predicted changes in relational reasoning for the 43 participants who completed a matrix reasoning task at two timepoints. Importantly, participants generally improved from timepoint 1 (mean ± sd = 23.3 ± 7.63) to timepoint 2 (mean ± sd = 27.2 ± 5.37; t = -5.01, *p* = .00001, *d* = -0.76, change mean ± sd = 3.9 ± 5.11). To relate the aforementioned changes in morphology to these changes in cognitive ability, we leveraged a previously-published, data-driven pipeline (Voorhies et al., 2021; Yao et al., 2022) to determine which of the LPFC sulci (Extended Data **Fig. 1-1**), if any, showed a change in cortical thickness (ΔCT) associated with this change in matrix reasoning score (ΔMR). To this end, we implemented a LASSO regression with cross-validation (Materials and Methods). A LASSO regression not only allows us to select sulci in a data-driven manner but also improves the generalizability of a model and prevents overfitting, especially in situations where there are 10 < x < 25 predictors (Heinze et al., 2018). We assessed the relationship between ΔCT and ΔMR separately in each hemisphere.

Implementing the LASSO regression revealed that 4 of the 17 left-hemisphere LPFC sulci in this study were selected at the α value that minimized cross-validated MSE (α = 0.07, MSE = 24.28; **Fig. 4A**), and thus associated with ΔMR. None of the LPFC sulci in the right hemisphere were selected (α = 0.5, MSE = 26.76). Specifically, the *pre-triangular sulcus* (prts; β *=* 2.89) and the *ascending ramus of the lateral fissure* (aalf; β *=* 14.9) showed positive relationships with ΔMR, whereas the *lateral marginal frontal sulcus* (lfms; β *=* -8.43) and the *anterior component of the posterior middle frontal sulcus* (pmfs-a; β *=* -1.49) showed negative relationships with ΔMR.

To further examine the relationship between the ΔCT of the selected LPFC sulci and ΔMR, we used the *LASSO-derived model* derived from our LASSO regression to predict ΔMR performance and conducted nested model comparisons to evaluate the fit of this LASSO-derived model (Materials and Methods). All models were fitted with LOOCV. The LASSO-derived model included the ΔCT of the left-hemisphere prts, aalf, lfms, and pmfs-a as predictors of ΔMR in the LOOCV linear regression. Baseline age was also included as a predictor in the model to assess whether this model outperformed baseline age alone. We found that the LASSO-derived model was associated with ΔMR (*R*^2^_CV_ = 0.22, MSE_CV_ = 19.94) and showed a moderate relationship between predicted and measured ΔMR (Spearman’s rho = 0.49, *p* = .0008; **Fig. 4B**).

When compared with a nested cross-validated model with all 17 LPFC sulci in the left hemisphere in this study and baseline age, we found that the addition of the other 13 sulci greatly weakened the model fit (*R*^2^_CV_ < 0.01, MSE_CV_ = 41.69; **Fig. 4C**). This comparison was consistent with the predictions of the LASSO regression. To determine whether the LASSO-derived model explained unique variance in ΔMR relative to baseline age, we compared this model to a nested cross-validated model with baseline age as the sole predictor. Compared to the baseline age model (*R*^2^_CV_ = 0.098, MSE_CV_ = 23.03; **Fig. 4C**), the LASSO-derived model showed increased prediction accuracy and decreased MSE_CV_ (**Fig. 4C**). Thus, the inclusion of the four selected LPFC sulci improved the prediction of ΔMR above and beyond baseline age; however, the addition of the remaining 13 LPFC sulci weakened the fit.

## Discussion

To our knowledge, this is the first study to examine how the morphology of specific LPFC sulci changes cross-sectionally and longitudinally, as well as whether a developmental relationship exists between LPFC sulcal morphology and cognitive performance. This work builds on the rich literature examining how cortical morphology changes during child development (e.g., Gogtay et al., 2004; Lebel et al., 2008; Shaw et al., 2008; Alemán-Gómez et al., 2013; Fjell et al., 2015; Amlien et al., 2016; Tamnes et al., 2017; Norbom et al., 2021, 2022; Baum et al., 2022; Bethlehem et al., 2022; Fuhrmann et al., 2022) and how LPFC maturation is linked to reasoning performance (e.g., Crone et al., 2009; Dumontheil et al., 2010; Wendelken et al., 2017; Leonard et al., 2019). Importantly, these findings reveal, at a more granular level, yet another way in which PFC is linked to reasoning performance (Milner and Petrides, 1984; Christoff et al., 2001; Vendetti and Bunge, 2014; Aichelburg et al., 2016; Urbanski et al., 2016; Hartogsveld et al., 2018; Assem et al., 2020; Holyoak and Monti, 2021). Leveraging one of the largest samplings of LPFC sulci within individual participants, we observed a striking complexity in how the morphology of a single cortical region changes. Further, our results show that longitudinal morphological changes in four left-hemisphere LPFC sulci are cognitively relevant during middle childhood and adolescence.

### On the development of sulcal morphology

We found that some, but not all, morphological features of LPFC sulci differ cross-sectionally between age groups and change longitudinally in children and adolescents. Specifically, while most morphological features differed between children/adolescents and young adults (i.e., depth, surface area, thickness, and *l*GI), only cortical thickness showed cross-sectional differences between children and adolescents and longitudinal changes during this time frame (6-18 years).

These age-related differences were not homogenous across our data-driven sulcal clusters or between cerebral hemispheres. This complexity suggests that a more nuanced approach is necessary to understand cortical development. As the field of neuroscience debates the importance of, and differences between, “large N” and “deep imaging” approaches (e.g., Genon et al., 2022; Gratton et al., 2022) our results and prior work (e.g., Raznahan et al., 2011; Alemán-Gómez et al., 2013; Cachia et al., 2016; Bethlehem et al., 2022; Fuhrmann et al., 2022; Willbrand et al., 2022a) emphasize that deep analysis of the kind that is most easily accomplished in smaller samples is crucial for delineating how, when, and where changes in cortical morphology occur during development.

Additionally, the lack of cross-sectional and longitudinal results regarding some morphological features (notably depth, *l*GI, and surface area) does not imply that they do not change during childhood and adolescence (Alemán-Gómez et al., 2013). Rather, we may not see cross-sectional differences during this age range if more consistent and/or more pronounced developmental changes take place earlier in child development, and the neural mechanisms leading to changes in the morphology of these individual sulci may occur over a longer period of time (> 1.56-year difference, on average) or a different time frame (6-18 years old) than what was studied here. Conversely, our results did present a difference in most features between children/adolescents and young adults. It is also plausible then that the most pronounced changes in LPFC sulcal morphology occur during the prenatal period and early childhood (Dubois et al., 2008; Meng et al., 2014; Le Guen et al., 2018; Im and Grant, 2019; Aslan Çetin and Madazlı, 2021), and again in older adulthood (Liu et al., 2013; Madan, 2021; Tang et al., 2021; Willbrand et al., 2022a).

### On the relationship between LPFC and reasoning

Using a data-driven approach we found that changes in the gray matter thickness of four left-hemisphere LPFC sulci predicted changes in reasoning performance. Strikingly, two of the four selected sulci, lfms and prts, likely border or fall within rostrolateral PFC (RLPFC)—a sub-region of LPFC implicated in reasoning based on both neuropsychological and fMRI data (Milner and Petrides, 1984; Christoff et al., 2001; Vendetti and Bunge, 2014; Aichelburg et al., 2016; Urbanski et al., 2016; Hartogsveld et al., 2018; Assem et al., 2020; Holyoak and Monti, 2021). Indeed, these two sulci co-localize, at the group level, with activation strongly linked to reasoning in a meta-analysis comparing 182 fMRI studies including this term with over 14,000 studies that did not include this term (tested via *Neurosynth*; **Fig. 5**). The other two implicated sulci, pmfs-a and aalf, appear to border dorsolateral and ventrolateral PFC regions that were also implicated in reasoning in the fMRI meta-analysis; of these, the dorsolateral sulcus was linked more closely to reasoning than to other cognitive processes.

**Figure 5.**
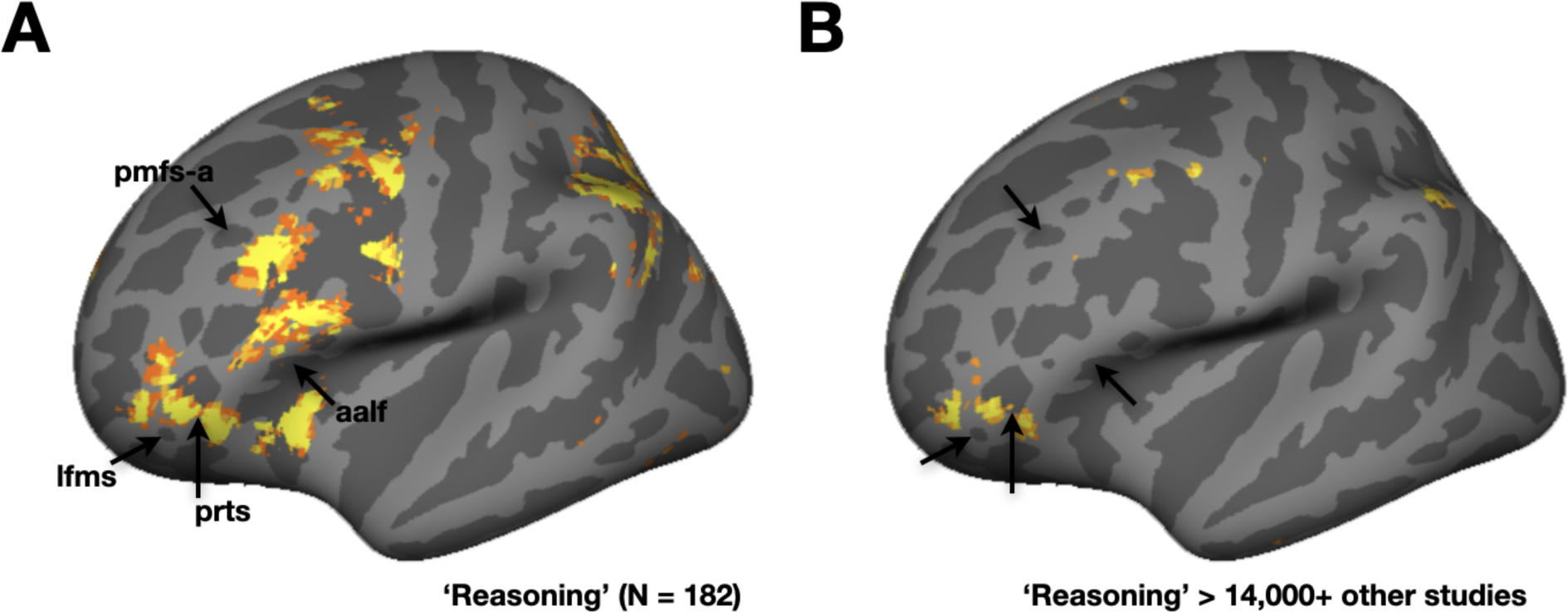
Selected LPFC sulci co-localize with functional activation for reasoning-related keywords in a meta-analysis using NeuroSynth. ***A***. Left hemisphere inflated *fsaverage* cortical surface displaying overlap visualization of a whole-brain FDR-corrected (*p <* 0.01) uniformity-test meta-analysis z-score map of the ‘reasoning’ term (downloaded from Neurosynth; https://neurosynth.org/). This map was generated from a Chi-squared test comparing the activation in each voxel for studies containing the term (N = 182) compared to what one would expect if activation were uniformly distributed throughout the gray matter. The four left-hemisphere LPFC sulci selected by the LASSO regression are identified on this surface with black arrows. ***B***. Same as *A*, but instead displaying overlap visualization of a whole-brain FDR-corrected (*p* < 0.01) association-test meta-analysis z-score map of the ‘reasoning’ term. This map was generated from a Chi-squared test comparing the proportion of studies demonstrating activation in each voxel for studies containing the term (N = 182) of interest compared to all other studies in the database (N > 14,000).

Building on prior work implicating general areas of LPFC to reasoning, our results suggest four small individually identified sulci that may serve as personalized “coordinates” of a larger cognitive globe (Stuss and Benson, 1984; see Miller et al., 2021a); integrating various findings, we could further understand the neuroanatomical substrates of reasoning at a more fine-grained level than previously considered, in concert with other anatomical (not to mention functional) features within and beyond LPFC subregions that have been linked to reasoning performance, both involving the current dataset (Wendelken et al., 2017; Voorhies et al., 2021; Willbrand et al., 2022b) and others (e.g., Ritchie et al., 2015; Chen et al., 2020). Other factors to consider are that anatomical relations to reasoning can vary over time (Shaw et al., 2006), and can depend on the population under consideration, with children from higher and lower socioeconomic backgrounds exhibiting different trajectories of cortical thinning during childhood (Piccolo et al., 2016) and different relations between cortical thickness and reasoning (Leonard et al., 2019).

### On relationships between sulcal morphology and human behavior

Sanides (1964) proposed a classic theory linking tertiary sulci, the last sulci to emerge in gestation, to the later-developing cognitive abilities supported by association cortices. His theory specifically focused on LPFC, wherein he proposed that the late emergence and continued postnatal morphological development of tertiary sulci are likely related to the cognitive skills associated with LPFC, which also show protracted development (Sanides, 1964). Our findings provide the first empirical support for the longitudinal aspect of this theory by showing that one such feature of specific LPFC tertiary sulci (gray matter thickness) is predictive of the development of one such ability (reasoning).

These findings also build upon prior research identifying that the *trajectory* of change in cortical thickness, not necessarily the thickness *itself*, is linked to reasoning (Shaw et al., 2006; Burgaleta et al., 2014; Schnack et al., 2015). However, these studies did not consider the cortex buried within *specific* cortical folds across individuals. Thus, this study extends this relationship to specific areas of cortex buried in sulci in LPFC. Mechanistically, cortical thinning likely reflects not only gray matter thinning but also the myelination of fibers extending into the cortical ribbon (Paus, 2005; de Faria et al., 2021; Norbom et al., 2021; Baum et al., 2022). Indeed, recent work by Natu and colleagues (2019) found that myelination is a key contributor to cortical thinning in the visual cortex during childhood. Therefore, changes in sulcal cortical thickness may reflect changes in the local cellular architecture and neural circuits that ultimately support the development of reasoning abilities during childhood and adolescence—a relationship that should be explored in future research.

Our findings also extend prior research on the relationship between sulcal morphology and behavior from cross-sectional individual differences to longitudinal changes within participants. For the most part, recent work has found that individual differences in sulcal depth, especially in LPFC, are predictive of individual differences in cognitive tasks during childhood and adolescence, more so than individual differences in sulcal thickness (Voorhies et al., 2021; Yao et al., 2022). However, as mentioned earlier in this Discussion, our findings and previous work (Shaw et al., 2006; Burgaleta et al., 2014; Schnack et al., 2015) emphasize that it is the trajectory of thinning that is cognitively meaningful and that depth does not appear to change during childhood and adolescence, both cross-sectionally and longitudinally. Nevertheless, sulcal depth could change over a longer period of time and a different time frame than what was studied here. Accordingly, future research should discern whether depth changes do occur in these sulci over a longer longitudinal period of time during middle childhood and adolescence and, if so, determine whether these changes are cognitively relevant. In closing, these results relay the importance of considering individual differences in neuroanatomy when studying the neurodevelopment of the cerebral cortex and lay the foundation for more precise structural-functional and structural-behavioral analyses in the future.

## Acknowledgments

This research was supported by NICHD R21HD100858 (Weiner, Bunge) and NSF CAREER Award 2042251 (Weiner). Funding for the original data collection and curation of the pediatric sample was provided by NINDS R01 NS057156 (Bunge, Ferrer) and NSF BCS1558585 (Bunge, Wendelken). We thank former members of the Bunge laboratory for assistance with data collection, and the families who participated in the original study. Data for the young adult sample were provided by the Human Connectome Project, WU-Minn Consortium (Principal Investigators: David Van Essen and Kamil Ugurbil; National Institutes of Health (NIH) Grant 1U54-MH-091657) funded by the 16 NIH Institutes and Centers that support the NIH Blueprint for Neuroscience Research and by the McDonnell Center for Systems Neuroscience at Washington University. We also thank Willa I. Voorhies, Jewelia K. Yao, Ishana Raghuram, and Jacob A. Miller for their assistance in defining the lateral prefrontal cortex sulci and developing the code used for this project.

