## Supplementary Materials for "Development of human lateral prefrontal sulcal morphology and its relation to reasoning performance"

### Extended Data

| No. | Abbreviation | Name |
|---|---|---|
| 1 | *cs* | central sulcus |
| 2 | *sprs* | superior precentral sulcus |
| 3 | *iprs* | inferior precentral sulcus |
| 4 | *ifs* | inferior frontal sulcus |
| 5 | *sfs-p* | superior frontal sulcus — posterior |
| 6 | *sfs-a* | superior frontal sulcus — anterior |
| 7 | *imfs-h* | intermediate frontal sulcus — horizontal |
| 8 | *imfs-v* | intermediate frontal sulcus — vertical |
| 9 | *pmfs-p* | posterior middle frontal sulcus — posterior |
| 10 | *pmfs-i* | posterior middle frontal sulcus — intermediate |
| 11 | *pmfs-a* | posterior middle frontal sulcus — anterior |
| 12 | *pimfs+* | paraintermediate frontal sulcus |
| 13 | *ds* | diagonal sulcus |
| 14 | *aalf* | ascending ramus of the lateral fissure |
| 15 | *ts* | triangular sulcus |
| 16 | *half* | horizontal ramus of the lateral fissure |
| 17 | *prts* | pretriangular sulcus |
| 18 | *lfms* | lateral frontomarginal sulcus |

**Figure 1-1. Sulcal definitions in lateral prefrontal cortex (LPFC).** Table of the 18 LPFC sulci explored in the present study. Numbers correspond to sulci in Figures 1 and 2. Sulcal abbreviations and the full name are included. +The pimfs can have 0, 1, or 2 components (dorsal and ventral; Voorhies et al., 2021; Yao et al., 2022; Willbrand et al., 2022). The pimfs is not included in our analyses due to this variability.

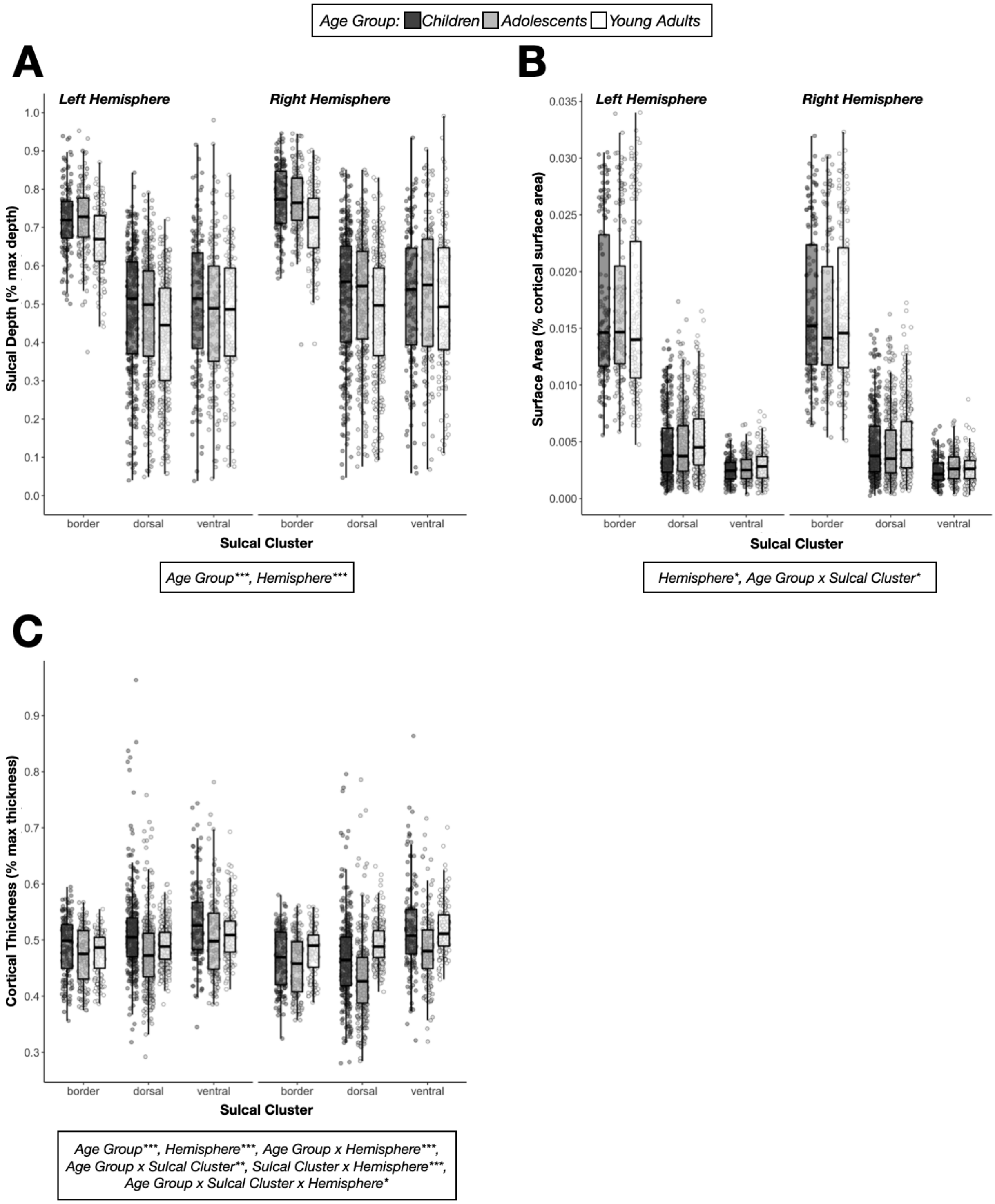


**Figure 3-1. Cross-sectional results with normalized morphological metrics.** ***A***. Same format as Figure 3 except for normalized sulcal depth (% max depth). Asterisks represent the same *p*-values ranges (**p* < .05; ***p* < .01; ****p*

< .001). ***B***. Same as *A,* but for normalized surface area (% cortical surface area). ***C***. Same as *A,* but for normalized mean cortical thickness (% max thickness).

| Effect | DF1 | DF2 | F-value | p-value | Significance | η2G |
|---|---|---|---|---|---|---|
| Raw (mm) | | | | | | |
| age_group | 2.000 | 102.000 | 30.102 | < 0.001 | *** | 0.165 |
| gender | 1.000 | 102.000 | 4.351 | 0.13 | n.s. | 0.014 |
| age_group:gender | 2.000 | 102.000 | 2.355 | 0.25 | n.s. | 0.015 |
| hemi | 1.000 | 102.000 | 26.216 | < 0.001 | *** | 0.029 |
| age_group:hemi | 2.000 | 102.000 | 0.607 | 0.684 | n.s. | 0.001 |
| gender:hemi | 1.000 | 102.000 | 1.415 | 0.401 | n.s. | 0.002 |
| age_group:gender:hemi | 2.000 | 102.000 | 0.021 | 0.988 | n.s. | 0.000 |
| age_group:sulcal_cluster | 3.703 | 188.849 | 2.377 | 0.174 | n.s. | 0.017 |
| gender:sulcal_cluster | 1.851 | 188.849 | 2.466 | 0.235 | n.s. | 0.009 |
| age_group:gender:sulcal_cluster | 3.703 | 188.849 | 1.765 | 0.305 | n.s. | 0.013 |
| hemi:sulcal_cluster | 1.643 | 167.630 | 1.725 | 0.344 | n.s. | 0.003 |
| age_group:hemi:sulcal_cluster | 3.287 | 167.630 | 1.747 | 0.306 | n.s. | 0.006 |
| gender:hemi:sulcal_cluster | 1.643 | 167.630 | 0.975 | 0.545 | n.s. | 0.002 |
| age_group:gender:hemi:sulcal_cluster | 3.287 | 167.630 | 0.165 | 0.96 | n.s. | 0.001 |
| Normalized (% max depth) | | | | | | |
| age_group | 2.000 | 102.000 | 13.947 | < 0.001 | *** | 0.097 |
| gender | 1.000 | 102.000 | 0.871 | 0.537 | n.s. | 0.003 |
| age_group:gender | 2.000 | 102.000 | 0.011 | 0.989 | n.s. | 0.000 |
| hemi | 1.000 | 102.000 | 85.057 | < 0.001 | *** | 0.111 |
| age_group:hemi | 2.000 | 102.000 | 0.638 | 0.671 | n.s. | 0.002 |
| gender:hemi | 1.000 | 102.000 | 0.816 | 0.545 | n.s. | 0.001 |
| age_group:gender:hemi | 2.000 | 102.000 | 0.105 | 0.936 | n.s. | 0.000 |
| age_group:sulcal_cluster | 3.651 | 186.225 | 2.198 | 0.213 | n.s. | 0.013 |
| gender:sulcal_cluster | 1.826 | 186.225 | 4.488 | 0.052 | n.s. | 0.013 |
| age_group:gender:sulcal_cluster | 3.651 | 186.225 | 1.416 | 0.401 | n.s. | 0.008 |
| hemi:sulcal_cluster | 1.663 | 169.639 | 2.052 | 0.305 | n.s. | 0.003 |
| age_group:hemi:sulcal_cluster | 3.326 | 169.639 | 1.770 | 0.305 | n.s. | 0.005 |
| gender:hemi:sulcal_cluster | 1.663 | 169.639 | 0.924 | 0.559 | n.s. | 0.001 |
| age_group:gender:hemi:sulcal_cluster | 3.326 | 169.639 | 0.149 | 0.962 | n.s. | 0.000 |

**Figure 3-2. Mixed model ANOVA results for sulcal depth.** All p-values are FDR corrected for multiple comparisons

(Significance: n.s. $p > .05$, $^{*}p < .05$, $^{**}p < .01$, $^{***}p < .001$). Effect size is reported as general eta squared (η2G). See Materials and Methods for additional analysis details.

| Effect | DF1 | DF2 | F-value | p-value | Significance | η2G |
|---|---|---|---|---|---|---|
| Raw (mm2) | | | | | | |
| age_group | 2.000 | 102.000 | 0.693 | 0.662 | n.s. | 0.005 |
| gender | 1.000 | 102.000 | 11.601 | 0.004 | ** | 0.038 |
| age_group:gender | 2.000 | 102.000 | 6.821 | 0.007 | ** | 0.045 |
| hemi | 1.000 | 102.000 | 3.338 | 0.2 | n.s. | 0.002 |
| age_group:hemi | 2.000 | 102.000 | 1.300 | 0.434 | n.s. | 0.001 |
| gender:hemi | 1.000 | 102.000 | 0.136 | 0.776 | n.s. | 0.000 |
| age_group:gender:hemi | 2.000 | 102.000 | 0.487 | 0.701 | n.s. | 0.000 |
| age_group:sulcal_cluster | 2.700 | 137.692 | 2.482 | 0.2 | n.s. | 0.021 |
| gender:sulcal_cluster | 1.350 | 137.692 | 1.943 | 0.313 | n.s. | 0.008 |
| age_group:gender:sulcal_cluster | 2.700 | 137.692 | 6.121 | 0.004 | ** | 0.050 |
| hemi:sulcal_cluster | 1.321 | 134.749 | 0.205 | 0.776 | n.s. | 0.000 |
| age_group:hemi:sulcal_cluster | 2.642 | 134.749 | 1.601 | 0.357 | n.s. | 0.005 |
| gender:hemi:sulcal_cluster | 1.321 | 134.749 | 1.379 | 0.417 | n.s. | 0.002 |
| age_group:gender:hemi:sulcal_cluster | 2.642 | 134.749 | 0.377 | 0.79 | n.s. | 0.001 |
| Normalized (% cortical surface area) | | | | | | |
| age_group | 2.000 | 102.000 | 0.668 | 0.662 | n.s. | 0.002 |
| gender | 1.000 | 102.000 | 2.598 | 0.269 | n.s. | 0.004 |
| age_group:gender | 2.000 | 102.000 | 0.859 | 0.61 | n.s. | 0.003 |
| hemi | 1.000 | 102.000 | 7.788 | 0.025 | * | 0.006 |
| age_group:hemi | 2.000 | 102.000 | 1.302 | 0.434 | n.s. | 0.002 |
| gender:hemi | 1.000 | 102.000 | 0.088 | 0.806 | n.s. | 0.000 |
| age_group:gender:hemi | 2.000 | 102.000 | 0.339 | 0.776 | n.s. | 0.000 |
| age_group:sulcal_cluster | 2.957 | 150.830 | 3.815 | 0.046 | * | 0.035 |
| gender:sulcal_cluster | 1.479 | 150.830 | 3.632 | 0.133 | n.s. | 0.017 |
| age_group:gender:sulcal_cluster | 2.957 | 150.830 | 1.979 | 0.287 | n.s. | 0.018 |
| hemi:sulcal_cluster | 1.316 | 134.185 | 0.603 | 0.657 | n.s. | 0.002 |
| age_group:hemi:sulcal_cluster | 2.631 | 134.185 | 1.580 | 0.36 | n.s. | 0.009 |
| gender:hemi:sulcal_cluster | 1.316 | 134.185 | 1.304 | 0.434 | n.s. | 0.004 |
| age_group:gender:hemi:sulcal_cluster | 2.631 | 134.185 | 0.409 | 0.776 | n.s. | 0.002 |

**Figure 3-3. Mixed model ANOVA results for surface area.** All p-values are FDR corrected for multiple

comparisons (Significance: n.s. $p > .05$, *$p < .05$, **$p < .01$, ***$p < .001$). Effect size is reported as general eta squared ($\eta 2G$). See Materials and Methods for additional analysis details. Although there was no *age group* main effect or interactions on raw surface area, there was an *age group* x *sulcal cluster* effect on normalized surface area. Here, the normalized surface area of dorsal sulci was smaller in children and adolescents than in young adults ($p$s < .0006, Tukey's adjustment; 9.28% and 9.48% less on average), but comparable between children and adolescents ($p$ = .99, Tukey's adjustment; Extended Data **Fig. 3-1**). The normalized surface area of ventral sulci was smaller in children than young adults ($p$ = .03, Tukey's adjustment; around 10.2% less on average), an effect driven largely by a marginal difference between children and adolescents ($p$ = .10, Tukey's adjustment), as there was not a significant difference between adolescents and young adults ($p$ = .99, Tukey's adjustment; Extended Data **Fig. 3-1**). Unlike the dorsal and ventral sulci, no age group differences were observed for border sulci ($p$s > .19, Tukey's adjustment; Extended Data **Fig. 3-1**).

| Effect | DF1 | DF2 | F-value | p-value | Significance | η2G |
|---|---|---|---|---|---|---|
| Raw (mm) | | | | | | |
| age_group | 2.000 | 102.000 | 14.808 | < 0.001 | *** | 0.118 |
| gender | 1.000 | 102.000 | 3.015 | 0.224 | n.s. | 0.013 |
| age_group:gender | 2.000 | 102.000 | 1.949 | 0.305 | n.s. | 0.017 |
| hemi | 1.000 | 102.000 | 58.332 | < 0.001 | *** | 0.076 |
| age_group:hemi | 2.000 | 102.000 | 24.132 | < 0.001 | *** | 0.063 |
| gender:hemi | 1.000 | 102.000 | 0.576 | 0.624 | n.s. | 0.001 |
| age_group:gender:hemi | 2.000 | 102.000 | 0.478 | 0.701 | n.s. | 0.001 |
| age_group:sulcal_cluster | 3.419 | 174.371 | 5.355 | 0.004 | ** | 0.023 |
| gender:sulcal_cluster | 1.710 | 174.371 | 1.757 | 0.344 | n.s. | 0.004 |
| age_group:gender:sulcal_cluster | 3.419 | 174.371 | 1.750 | 0.305 | n.s. | 0.008 |
| hemi:sulcal_cluster | 1.602 | 163.367 | 16.373 | < 0.001 | *** | 0.027 |
| age_group:hemi:sulcal_cluster | 3.203 | 163.367 | 3.609 | 0.047 | * | 0.012 |
| gender:hemi:sulcal_cluster | 1.602 | 163.367 | 0.599 | 0.662 | n.s. | 0.001 |
| age_group:gender:hemi:sulcal_cluster | 3.203 | 163.367 | 0.612 | 0.701 | n.s. | 0.002 |
| Normalized (% max thickness) | | | | | | |
| age_group | 2.000 | 102.000 | 14.835 | < 0.001 | *** | 0.118 |
| gender | 1.000 | 102.000 | 3.034 | 0.224 | n.s. | 0.013 |
| age_group:gender | 2.000 | 102.000 | 1.957 | 0.305 | n.s. | 0.017 |
| hemi | 1.000 | 102.000 | 58.511 | < 0.001 | *** | 0.076 |
| age_group:hemi | 2.000 | 102.000 | 24.243 | < 0.001 | *** | 0.064 |
| gender:hemi | 1.000 | 102.000 | 0.570 | 0.624 | n.s. | 0.001 |
| age_group:gender:hemi | 2.000 | 102.000 | 0.484 | 0.701 | n.s. | 0.001 |
| age_group:sulcal_cluster | 3.413 | 174.052 | 5.348 | 0.004 | ** | 0.023 |
| gender:sulcal_cluster | 1.706 | 174.052 | 1.736 | 0.344 | n.s. | 0.004 |
| age_group:gender:sulcal_cluster | 3.413 | 174.052 | 1.756 | 0.305 | n.s. | 0.008 |
| hemi:sulcal_cluster | 1.599 | 163.117 | 16.419 | < 0.001 | *** | 0.027 |
| age_group:hemi:sulcal_cluster | 3.198 | 163.117 | 3.605 | 0.047 | * | 0.012 |
| gender:hemi:sulcal_cluster | 1.599 | 163.117 | 0.594 | 0.662 | n.s. | 0.001 |
| age_group:gender:hemi:sulcal_cluster | 3.198 | 163.117 | 0.609 | 0.701 | n.s. | 0.002 |

**Figure 3-4. Mixed model ANOVA results for cortical thickness.** All p-values are FDR corrected for multiple

comparisons (Significance: n.s. $p > .05$, *$p < .05$, **$p < .01$, ***$p < .001$). Effect size is reported as general eta squared ($\eta 2G$). See Materials and Methods for additional analysis details. See Extended Data Fig. 3-6 for post hoc results on the 3-way interaction.

| Effect | DF1 | DF2 | F-value | p-value | Significance | η2G |
|---|---|---|---|---|---|---|
| age_group | 2.000 | 102.000 | 29.877 | < 0.001 | *** | 0.254 |
| gender | 1.000 | 102.000 | 4.122 | 0.139 | n.s. | 0.023 |
| age_group:gender | 2.000 | 102.000 | 0.563 | 0.697 | n.s. | 0.006 |
| hemi | 1.000 | 102.000 | 5.939 | 0.056 | n.s. | 0.006 |
| age_group:hemi | 2.000 | 102.000 | 0.573 | 0.697 | n.s. | 0.001 |
| gender:hemi | 1.000 | 102.000 | 1.234 | 0.434 | n.s. | 0.001 |
| age_group:gender:hemi | 2.000 | 102.000 | 1.957 | 0.305 | n.s. | 0.004 |
| age_group:sulcal_cluster | 2.696 | 137.505 | 0.793 | 0.657 | n.s. | 0.003 |
| gender:sulcal_cluster | 1.348 | 137.505 | 0.738 | 0.61 | n.s. | 0.002 |
| age_group:gender:sulcal_cluster | 2.696 | 137.505 | 0.591 | 0.701 | n.s. | 0.002 |
| hemi:sulcal_cluster | 1.385 | 141.295 | 1.572 | 0.376 | n.s. | 0.002 |
| age_group:hemi:sulcal_cluster | 2.770 | 141.295 | 1.219 | 0.47 | n.s. | 0.003 |
| gender:hemi:sulcal_cluster | 1.385 | 141.295 | 0.359 | 0.701 | n.s. | 0.000 |
| age_group:gender:hemi:sulcal_cluster | 2.770 | 141.295 | 0.419 | 0.776 | n.s. | 0.001 |

**Figure 3-5. Mixed model ANOVA results for local gyrification index.** All p-values are FDR corrected for multiple comparisons (Significance: n.s. $p > .05$, *$p < .05$, **$p < .01$, ***$p < .001$). Effect size is reported as general eta squared ($\eta 2G$). See Materials and Methods for additional analysis details.

| Hemisphere | ch - ad | p-value | ch - ya | p-value | ad - ya | p-value |
|---|---|---|---|---|---|---|
| Border sulci | | | | | | |
| left | 0.08 | .004 | 0.06 | .03 | -0.02 | .74 |
| right | 0.05 | .16 | -0.07 | .02 | -0.11 | .0001 |
| Dorsal sulci | | | | | | |
| left | 0.17 | .0001 | 0.11 | .005 | -0.05 | .37 |
| right | 0.16 | <.0001 | -0.13 | .0004 | -0.29 | <.0001 |
| Ventral sulci | | | | | | |
| left | 0.11 | .03 | 0.10 | .06 | -0.02 | .93 |
| right | 0.15 | .002 | -0.001 | .99 | -0.15 | .002 |

**Figure 3-6. Post-hoc comparisons for the age group x sulcal cluster x hemisphere interaction on cortical thickness (raw).** Post hoc pairwise comparisons (Tukey's Method) for the 3-way interaction between age group, sulcal cluster, and hemisphere on cortical thickness (mm; Extended Data Fig. 3-4). Shown here are the average differences between age groups for each sulcal cluster (border, dorsal, ventral) in each hemisphere (left, right) along with the associated adjusted p-values (Tukey test). Abbreviations are as follows: children (ch); adolescents (ad); young adults (ya). Border sulci showed a decline in cortical thickness from children to adolescents and young adults in the left hemisphere, whereas, in the right hemisphere, these sulci showed an increase in cortical thickness from children and adolescents to young adults. Dorsal sulci showed a similar relationship in the left hemisphere to border sulci, but in the right hemisphere, dorsal sulci showed a U-shaped function between age groups on cortical thickness. Ventral sulci showed slightly different U-shaped functions between age groups on cortical thickness in both hemispheres.

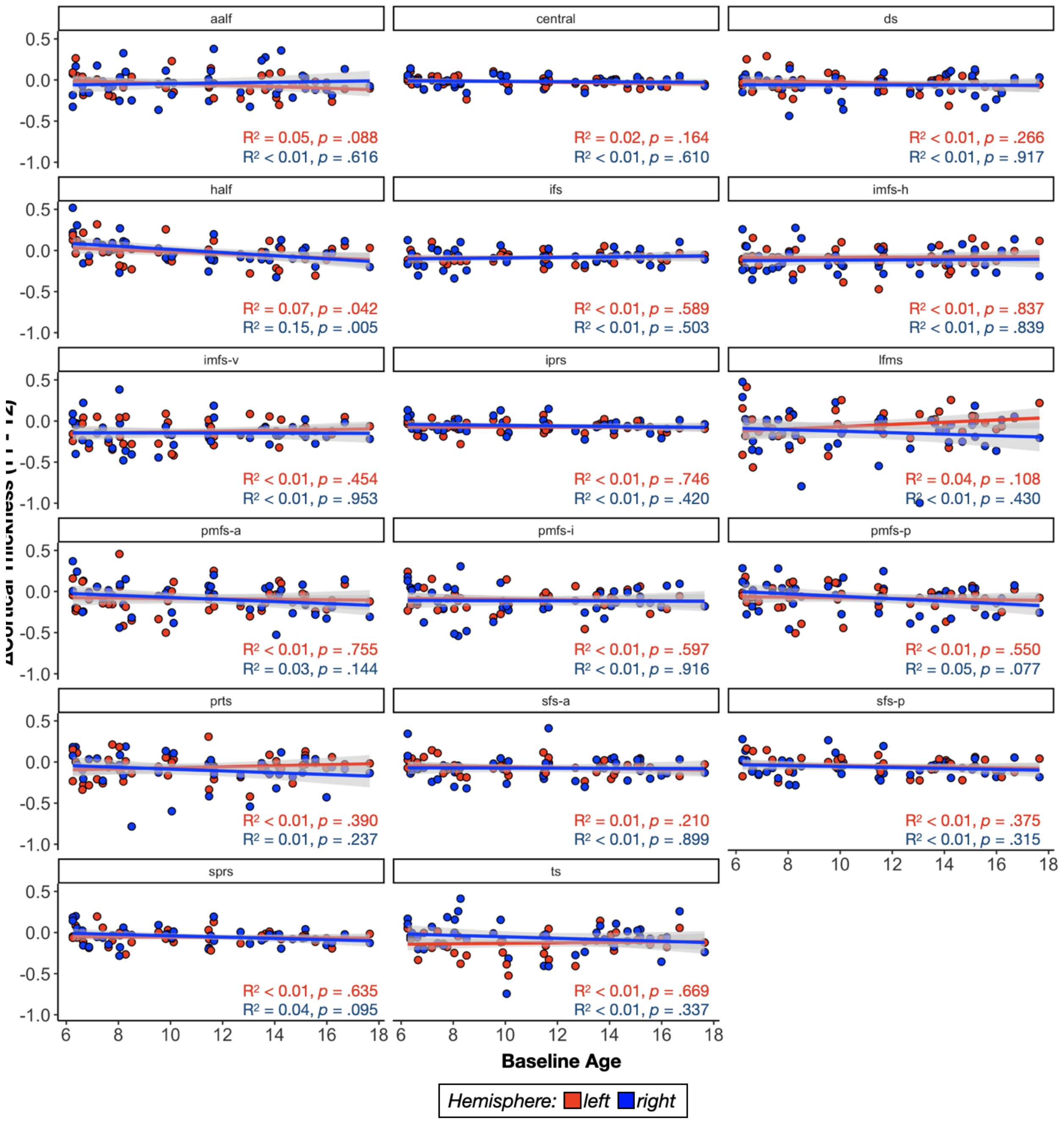


**Figure 4-1. Relationship between change in cortical thickness of each sulcus and baseline age.** Each subplot visualizes the change in cortical thickness of each of the 17 consistent LPFC sulci in the left (red) and right (blue) hemispheres as a function of baseline age. Individual dots represent individual participants and are colored by hemisphere (see the key below plots). The best fit line and the 95% confidence interval for each regression are also included. The $R^2$ and *p* values are reported at the bottom right of each subplot.

| | Count |
|---|---|
| Racial categories | |
| American Indian/Alaskan Native | 0 |
| Asian/Native Hawaiian/Other Pacific Islander | 5 |
| Black or African American | 4 |
| White | 47 |
| More Than One Race | 14 |
| Unknown or Not Reported | 2 |
| Ethnic categories | |
| Hispanic or Latino | 11 |
| Not Hispanic or Latino | 59 |
| Unknown or Not Reported | 2 |
| Highest degree earned by parent/guardian | |
| High School/GED | 8 |
| Vocational | 1 |
| Associate degree | 6 |
| Bachelor's degree | 18 |
| Master's degree | 15 |
| Doctorate | 4 |
| Professional | 3 |
| Other | 3 |
| None of the above (less than high school) | 1 |
| Unknown or Not Reported | 13 |
| Total household income | |
| $16,000-$24,999 | 2 |
| $25,000-$34,999 | 3 |
| $50,000-$74,999 | 6 |
| $75,000-$99,999 | 8 |
| $100,000-$199,999 | 27 |
| Over $200,000 | 5 |
| Unknown or Not Reported | 21 |

**Table 1-1. Demographic and socioeconomic information of the pediatric sample.** N = 72. All information is

parent/guardian reported.

| | Count |
|---|---|
| Racial categories | |
| American Indian/Alaskan Native | 1 |
| Asian/Native Hawaiian/Other Pacific Islander | 1 |
| Black or African American | 7 |
| White | 26 |
| Unknown or Not Reported | 1 |
| Ethnic categories | |
| Hispanic or Latino | 2 |
| Not Hispanic or Latino | 33 |
| Unknown or Not Reported | 1 |
| Years of education completed | |
| <11 | 1 |
| 12 | 9 |
| 13 | 2 |
| 14 | 2 |
| 15 | 2 |
| 16 | 13 |
| <17 | 7 |
| Total household income | |
| <$10,000 | 6 |
| $10,000-$19,999 | 5 |
| $20,000-$29,999 | 4 |
| $30,000-$39,999 | 4 |
| $40,000-$49,999 | 2 |
| $50,000-$74,999 | 4 |
| $75,000-$99,999 | 6 |
| Over $100,000 | 5 |

**Table 1-2. Demographic and socioeconomic information of the young adult sample.** N = 36.
